# Trends of Hospitalization in Acute Pancreatitis in Patients in the United States from 2001-2014

**DOI:** 10.1101/547273

**Authors:** Roberto Argo, Albert Bianco, Kevin Casey, Francesco Ricci, Daniele Costa

**Affiliations:** Department of Medicine, University of Verona, Verona, Italy; Department of Medical Sciences, University of Bologna, Bologna, Italy

**Author notes:** Correspondence, Department of Medicine, University of Verona, Verona, Italy.

**Keywords:** pancreatitis, epidemiology, etiology, risk factors, gallstones, alcohol

## Abstract

**Background & Purpose:** The prevalence of acute pancreatitis(AP) has increased over time and is one of the most important gastrointestinal causes of frequent admissions to hospital in the United States. The cost burden of AP has been steadily increasing. The primary objective of our study was to analyze patient demographics, cost burden, mortality and length of stay associated with AP hospital admissions.

**Methods:** Nationwide inpatient sample (NIS) database was used to identify AP admissions in all patients from ≥18 years of age from 2001 to 2014 using ICD-9-CM code 577.0 as the principal discharge diagnosis

**Results:** The number of hospitalizations increased from 215,238 in 2001 to 279,145 in 2014. Inhospital mortality decreased from 1.74% in 2001 to 0.66% in 2014. Mean length of hospital stay has decreased from 6.1 days to 4.6 days during the same period, but the mean hospital charges increased from $19,303 in 2001 to $35,728 in 2014. The proportion of males to females with acute pancreatitis is slowly trending up from 2001 to 2014.

**Conclusion:** The number of hospitalizations due to acute pancreatitis has been steadily increasing, and further research needs to be done on finding out the reasons for increased causes of hospitalization and ways to decrease the cost burden on patients and hospitals.

## 1. Introduction

Acute pancreatitis (AP) is an inflammatory condition of the pancreas which is characterized by abdominal pain and elevated pancreatic enzymes in the blood.(1) AP may range from a mild self-limiting form to a more severe form (acute necrotizing pancreatitis). Gallstones and chronic alcoholism are the two most common etiological factors associated with AP. Other causes include drugs, infections, hyperlipidemia, trauma, hypercalcemia, HIV, neoplasms and idiopathic.(2)

The prevalence of acute pancreatitis (AP) has increased over time and is one of the most important gastrointestinal causes of frequent admissions to hospital in the United States. The cost burden of AP has been steadily increasing. The primary objective of our study was to analyze patient demographics, cost burden, mortality and length of stay associated with AP hospital admissions.

## 2. Materials and Methods

Nationwide inpatient sample (NIS) database was used to identify AP admissions in all patients from >18 years of age from 2001 to 2014 using ICD-9-CM code 577.0 as the principal discharge diagnosis. NIS is the largest all-payer inpatient care database in the United States, containing data on more than 7 million hospital stays. It has a large sample size which is ideal for developing national and regional estimates.

We a carried a literature search for etiology of acute pancreatitis using the online databases of PubMed, Embase, Scopus, Google Scholar. We reviewed the pertinent articles about acute pancreatitis and its etiology. The review is an attempt to provide a better understanding of the possible various etiologic causes of acute pancreatitis.

## 3. Results

A total of 481,231,240 discharges with a diagnosis of AP were analyzed from 2001 to 2014 from the NIS database. Table 1 summarizes the data analyzed. The number of hospitalizations increased from 215,238 in 2001 to 279,145 in 2014. In-hospital mortality decreased from 1.74% in 2001 to 0.66% in 2014 (Figure 1). Mean length of hospital stay has decreased from 6.1 days to 4.6 days during the same period, but the mean hospital charges increased from $19,303 in 2001 to $35,728 in 2014. The proportion of males to females with acute pancreatitis is slowly trending up from 2001 to 2014. Aggregate charges increased from 4.15 billion dollars to 9.98 billion dollars.

**Figure 1.**
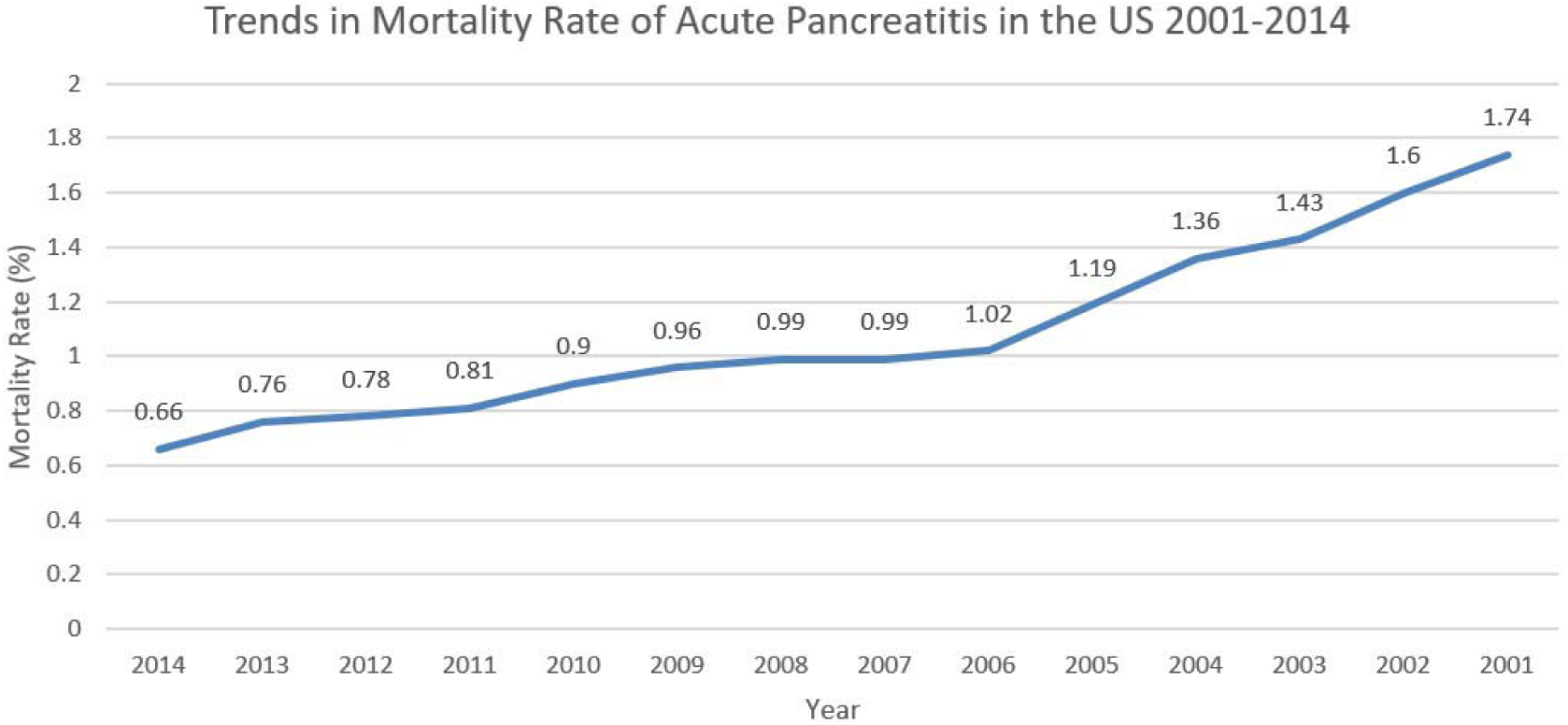
Trends in Mortality Rate of Acute Pancreatitis in the United States from 2001-2014.

**Table 1.**
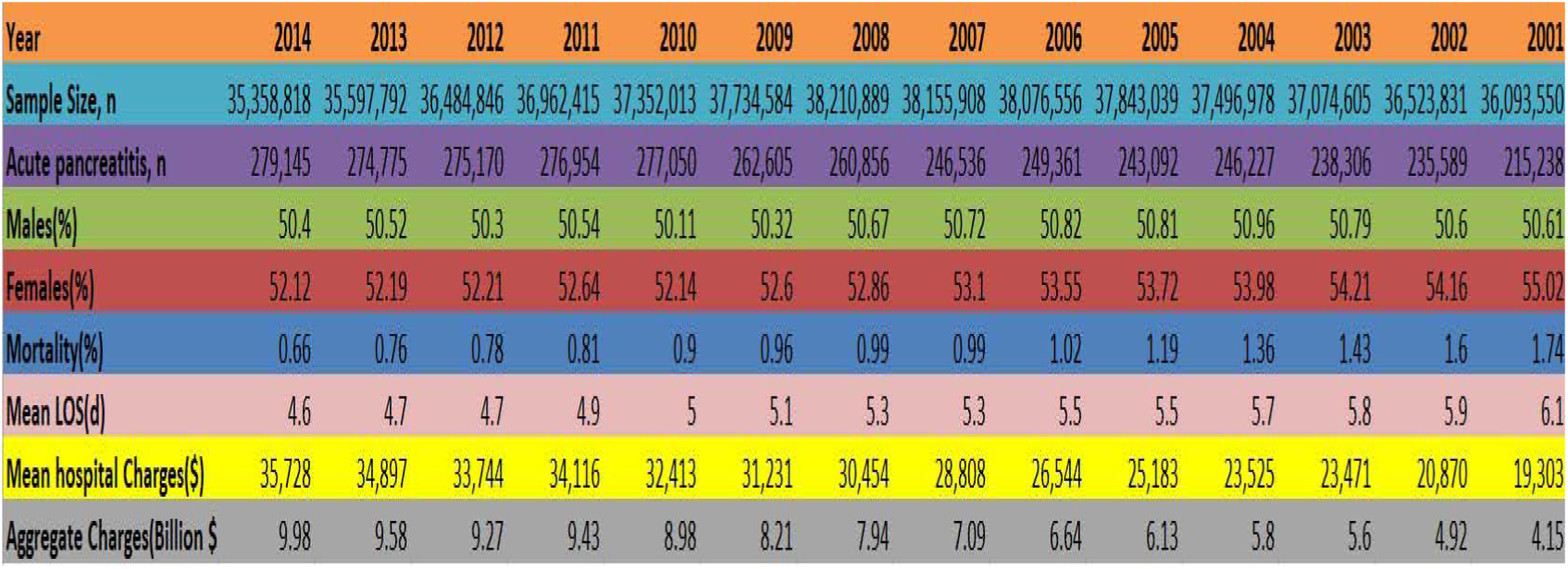
Trends of Hospitalization in Acute Pancreatitis in Patients in the United States from 2001-2014

| Year | 2014 | 2013 | 2012 | 2011 | 2010 | 2009 | 2008 | 2007 | 2006 | 2005 | 2004 | 2003 | 2002 | 2001 |
| --- | --- | --- | --- | --- | --- | --- | --- | --- | --- | --- | --- | --- | --- | --- |
| Sample Size, n | 35,358,818 | 35,597,792 | 36,484,846 | 36,962,415 | 37,352,013 | 37,734,584 | 38,210,889 | 38,155,908 | 38,076,556 | 37,843,039 | 37,496,978 | 37,074,605 | 36,523,831 | 36,093,550 |
| Acute pancreatitis, n | 279,145 | 274,775 | 275,170 | 276,954 | 277,050 | 262,605 | 260,856 | 246,536 | 249,361 | 243,092 | 246,227 | 238,306 | 235,589 | 215,238 |
| Males(%) | 50.4 | 50.52 | 50.3 | 50.54 | 50.11 | 50.32 | 50.67 | 50.72 | 50.82 | 50.81 | 50.96 | 50.79 | 50.6 | 50.61 |
| Females(%) | 52.12 | 52.19 | 52.21 | 52.64 | 52.14 | 52.6 | 52.86 | 53.1 | 53.55 | 53.72 | 53.98 | 54.21 | 54.16 | 55.02 |
| Mortality(%) | 0.66 | 0.76 | 0.78 | 0.81 | 0.9 | 0.96 | 0.99 | 0.99 | 1.02 | 1.19 | 1.36 | 1.43 | 1.6 | 1.74 |
| Mean LOS(d) | 4.6 | 4.7 | 4.7 | 4.9 | 5 | 5.1 | 5.3 | 5.3 | 5.5 | 5.5 | 5.7 | 5.8 | 5.9 | 6.1 |
| Mean hospital Charges(\$) | 35,728 | 34,897 | 33,744 | 34,116 | 32,413 | 31,231 | 30,454 | 28,808 | 26,544 | 25,183 | 23,525 | 23,471 | 20,870 | 19,303 |
| Aggregate Charges(Billion \$) | 9.98 | 9.58 | 9.27 | 9.43 | 8.98 | 8.21 | 7.94 | 7.09 | 6.64 | 6.13 | 5.8 | 5.6 | 4.92 | 4.15 |

| Year | 2014 | 2013 | 2012 | 2011 | 2010 | 2009 | 2008 | 2007 | 2006 | 2005 | 2004 | 2003 | 2002 | 2001 |
| --- | --- | --- | --- | --- | --- | --- | --- | --- | --- | --- | --- | --- | --- | --- |
| Sample Size, n | 35,358,818 | 35,597,792 | 36,484,846 | 36,962,415 | 37,352,013 | 37,734,584 | 38,210,889 | 38,155,908 | 38,076,556 | 37,843,039 | 37,496,978 | 37,074,605 | 36,523,831 | 36,093,550 |
| Acute pancreatitis, n | 279,145 | 274,775 | 275,170 | 276,954 | 277,050 | 262,605 | 260,856 | 246,536 | 249,361 | 243,092 | 246,227 | 238,306 | 235,589 | 215,238 |
| Males(%) | 50.4 | 50.52 | 50.3 | 50.54 | 50.11 | 50.32 | 50.67 | 50.72 | 50.82 | 50.81 | 50.96 | 50.79 | 50.6 | 50.61 |
| Females(%) | 52.12 | 52.19 | 52.21 | 52.64 | 52.14 | 52.6 | 52.86 | 53.1 | 53.55 | 53.72 | 53.98 | 54.21 | 54.16 | 55.02 |
| Mortality(%) | 0.66 | 0.76 | 0.78 | 0.81 | 0.9 | 0.96 | 0.99 | 0.99 | 1.02 | 1.19 | 1.36 | 1.43 | 1.6 | 1.74 |
| Mean LOS(d) | 4.6 | 4.7 | 4.7 | 4.9 | 5 | 5.1 | 5.3 | 5.3 | 5.5 | 5.5 | 5.7 | 5.8 | 5.9 | 6.1 |
| Mean hospital Charges(\$) | 35,728 | 34,897 | 33,744 | 34,116 | 32,413 | 31,231 | 30,454 | 28,808 | 26,544 | 25,183 | 23,525 | 23,471 | 20,870 | 19,303 |
| Aggregate Charges(Billion \$ | 9.98 | 9.58 | 9.27 | 9.43 | 8.98 | 8.21 | 7.94 | 7.09 | 6.64 | 6.13 | 5.8 | 5.6 | 4.92 | 4.15 |

## 4. Discussion

Gallstones are the most common cause of AP accounting for about 40 to 70 percent of cases.(3, 4). The risk of developing AP in patients with gallstones is more significant in men. However, women have a higher prevalence of gallstones, and the incidence of gallstone pancreatitis is higher in women. Patients with gallstones of size less than 5 mm are thought to be at higher risk of gallstone pancreatitis.(5)

Alcohol intake is responsible for approximately 25 to 35 percent of cases of AP in the United States.(6) Cigarette smoking might have an additive effect along with alcohol in inducing pancreatitis. Alcohol may increase the secretion of lysosomal enzymes by acting on the pancreatic acinar cells.(7) Although the exact pathogenesis of alcohol-induced AP is still unclear.(8)

Serum triglyceride levels above 1000 mg/dL can cause AP.(9–11) Hypertriglyceridemia accounts for about 1 to 14 percent of cases of AP.(12) Pancreatitis secondary to hypertriglyceridemia is usually seen in the presence of one or more factors such as diabetes, alcoholism in the presence of an underlying common genetic abnormality like familial hypertriglyceridemia.(12, 13) Insulin and heparin along with other measures used for AP have been proposed as the treatment for AP due to hypertriglyceridemia.(11)

Post-endoscopic retrograde cholangiopancreatography (ERCP) is associated with 3-5% of cases with AP.(14) The risk of Post ERCP AP is also related to the type of ERCP procedure being performed.(15) Studies have shown that patients undergoing ERCP for sphincter of Oddi dysfunction are more to develop AP when compared to other ERCP related procedures.(16)

Pancreatitis has been associated with multiple infections like Viruses (Mumps, hepatitis B, cytomegalovirus, herpes simplex, human immunodeficiency virus), Bacteria (Mycoplasma, Legionella, Salmonella), Fungi (Aspergillus) and Parasites (Toxoplasma, Ascaris).(17, 18)

Mutations in CTRC gene, CFTR gene, PRSS1 gene, SPINK1 have been implicated in pancreatitis etiology.(19) Patients with pancreatitis due to genetic mutations usually present as recurrent bouts of AP or childhood pancreatitis before it progresses to chronic pancreatitis and complications related to pancreatitis.

Drug-induced pancreatitis is rare and accounts for less than 5% of the cases.(20) Prognosis of drug-induced AP is good, and mortality is low. The diagnosis of drug-induced pancreatitis is difficult to establish as it is a diagnosis of exclusion.(21) Common drugs implicated in causing AP include aminosalicylates, sulfonamides, diuretics, valproic acid, didanosine, pentamidine, tetracycline, azathioprine, steroids.(22–24) Recently Glucagonlike peptide 1-based therapies have been increasingly implicated in etiology of drug-induced AP.(25)

Idiopathic causes can account up to 25% of the causes of AP. No etiology could be found on history, diagnostic tests or imaging. Research has shown recently that patient with idiopathic pancreatitis usually has complex genetic profiles.(26)

Other rare causes of pancreatitis include Pancreatic duct injury, Biliary obstruction, Hypercalcemia, Vascular disease, Biliary sludge and microlithiasis, Anatomic pancreatic anomalies.(27–33)

## 5. Conclusions

Etiology of AP can be established in most cases of AP. Although some cases can be challenging and in some patients, we may not find any etiology (idiopathic). Understanding the etiologic and risk factors of AP helps in formulating an appropriate management plan and helps to decrease the length of stay and cost associated with hospitalizations due to AP and might also lead to decrease in mortality due to AP. Overall in-hospital mortality decreased but the number of hospitalizations and mean cost has increased over the years. This represents a significant cost burden on the healthcare system and patients. The decrease in mortality can be attributed to early diagnosis and early institution of aggressive treatments. The number of hospitalizations due to acute pancreatitis has been steadily increasing, and further research needs to be done on finding out the reasons for increased causes of hospitalization and ways to decrease the cost burden on patients and hospitals.

## Author Contributions

Conception and design: Roberto Argo, Albert Bianco, Kevin Casey, Francesco Ricci, Daniele Costa.

Analysis and interpretation, Drafting and Critical revision of the Article: Roberto Argo, Albert Bianco, Kevin Casey, Francesco Ricci, Daniele Costa.

Final approval of the article: Roberto Argo, Albert Bianco, Kevin Casey, Francesco Ricci, Daniele Costa.

## Acknowledgments

None

